# Persistent activity in primate auditory cortex evoked by sensory stimulation

**DOI:** 10.1101/387837

**Authors:** James E. Cooke, Julie J. Lee, Edward L. Bartlett, Xiaoqin Wang, Daniel Bendor

## Abstract

Persistent activity, the elevated firing of a neuron after the termination of a stimulus, is hypothesized to play a critical role in working memory. This form of activity is therefore typically studied within the context of a behavioural task that includes a working memory component. Here we investigated whether persistent activity is observed in sensory cortex and thalamus in the absence of any explicit behavioural task. We recorded spiking activity from single units in the auditory cortex (fields A1, R and RT) and thalamus of awake, passively-listening marmosets. We observed persistent activity that lasted for hundreds of milliseconds following the termination of the acoustic stimulus, in the absence of a task. Persistent activity was observed following both adapting and sustained responses during the stimulus and showed similar stimulus tuning to these evoked responses. Persistent activity was also observed following suppression in firing during the stimulus. These response types were observed across all cortical fields tested, but were largely absent from thalamus. As well as being of shorter duration, thalamic persistent activity emerged following a longer latency than in cortex, indicating that persistent activity may be generated within auditory cortex during passive listening. Given that these responses were observed in the absence of a explicit behavioural task, persistent activity in sensory cortex may have functional importance beyond storing task-relevant information in working memory.

## Introduction

When an animal is required to hold information in working memory [1], neurons in a number of cortical areas have been found to fire persistently for durations on the order of seconds [2, 3]. In traditional working memory tasks, subjects are presented with a cue and are required to respond after a delay period, during which the cue is no longer available. Neurons in the dorsolateral prefrontal cortex of macaque have been found to fire persistently during such delay periods, providing a possible substrate for the maintenance of information in working memory [4, 2, 3, 5, 6, 7, 8]. In keeping with this proposed function, suppressing this activity has been shown to impair performance on working memory tasks [9, 10, 11].

Such activity has been found to carry spatial information such as the location of visual stimulus [12, 2, 13, 14] as well as the direction of a forthcoming saccade [8]. This activity has also been found to be involved in the retention of non-spatial information, such as the color or shape of an object [15, 15, 16]. In addition to prefrontal activity, sensory areas have been found to encode stimulus information across a delay period in the form of persistent activity. In a tone discrimination task where the tones were separated by a one-second delay period, neurons in the auditory cortex of the macaque were found to elevate their firing during the delay period [17]. Persistent responses have also been reported in human auditory cortex, as measured by magnetoencephalography (MEG) [18]. These findings demonstrate the ability of auditory cortical neurons to maintain their firing for hundreds of milliseconds in the absence of a coincident stimulus.

During passive listening, the termination of an auditory stimulus routinely evokes transient offset responses in auditory cortical neurons, typically lasting up to tens of milliseconds [19, 20, 21, 22]. In the cat, the tonal receptive fields of offset responses are often similar to the tuning of responses evoked at the onset of the stimulus although onset and offset receptive fields can also vary their tuning properties [21]. In the mouse however, the tuning of onset and offset responses is largely distinct [20]. In keeping with this inter-species variation in the tuning of offset responses, the mechanistic contribution of inhibition appears to vary across species. There is evidence from the cat and rat that offset responses may be due largely to rebound following inhibition from sensory stimulation [22, 21] while in the mouse this mechanism appears to contribute little to offset responses [20]. Despite the lack of consensus surrounding inhibition as a mechanism for generating offset responses, auditory cortical networks across all of these species have been found to routinely generate activity following the termination of effective auditory stimulus. The maximum duration of activity that can be evoked following stimulus offset, however, is currently unknown.

Here, we investigated whether it is possible to evoke long-duration post-stimulus activity in auditory cortical neurons in the absence of a behavioural task. We recorded activity from single units in the auditory cortex (fields A1, R and RT) and thalamus of awake, passively-listening marmosets while we presented an array of auditory stimuli [23, 24, 25, 26]. We observed persistent activity lasting for hundreds of milliseconds following the termination of the acoustic stimulus in a sub-population of auditory cortical neurons. This activity followed a variety of response profiles during sensory stimulation, including adapting, sustained and suppressed responses. Post-stimulus activity had shorter latency and was of longer duration in cortex than in thalamus, indicating that the mechanisms underlying this activity may be primarily cortical.

## Materials and methods

The data analysed in this study were obtained from several previous experiments conducted in the Laboratory of Auditory Neurophysiology at Johns Hopkins University School of Medicine [23, 24, 25, 26]. The methods to record single-unit activity in awake marmosets were previously described [27] and are briefly summarised below.

Single-unit recordings from auditory cortex and thalamus were conducted in awake, passively listening marmosets sitting on a semirestraint device with their head immobilized, within a double-walled soundproof chamber (Industrial Acoustics). The inside wall of the chamber was covered by 3-inch acoustic absorption foam (Sonex). For auditory cortical recordings high-impedance tungsten microelectrodes (3–5 M, A-M Systems) were inserted perpendicular to the cortical surface. Electrodes were mounted on a micromanipulator (Narishige) and advanced by a manual hydraulic microdrive (Trent Wells). Action potentials were detected on-line using a template-based spike sorter (Multi-Spike Detector; Alpha Omega Engineering) and continuously monitored by the experimenter while data recording progressed. Typically 5-15 electrode penetrations were made within a miniature recording hole (diameter ~1 mm) over the course of several days, after which the hole was sealed with dental cement and another hole opened for new electrode penetrations. Neurons were recorded from all cortical layers, but most commonly from supragranular layers.

### Generation of acoustic stimuli

Acoustic stimuli were generated digitally and delivered by a free-field loudspeaker located one meter directly in front of the animal. All sound stimuli were generated at a 100 kHz sampling rate and low-pass filtered at 50 kHz. The sound level of individual frequency components used in this study was no higher than 80 dB SPL. Frequency tuning curves and rate-level functions were generated using pure-tone stimuli of 200 ms in duration with interstimulus intervals of 500 ms, and had a minimum of five repetitions each. Stimulus presentation order was fully randomised. Pure-tone stimuli intensity levels were generally 10-20 dB above threshold for neurons with monotonic rate-level functions, or at preferred levels for non-monotonic neurons. Broadband rectangular clicks or narrowband clicks made of brief pulses of white noise or a tone (at an integer multiple of the frequency) were used to generate click trains. Rectangular click trains had a width of 0.1 ms while narrowband clicks had each pulse convolved with a Gaussian envelope with a standard deviation of 0.1-0.4.

### Identification of cortical fields

Single units with significant neuronal discharges to narrowband stimuli, such as tones and band-pass noise, were used to generate cortical characteristic frequency maps. The characteristic frequency of each location on the map was determined by the median characteristic frequency of all electrode tracks within 0.25 mm. Electrode track characteristic frequencies were calculated by computing the median characteristic frequency of units within the track. The anterior-posterior position is reported relative to the boundary between A1 and R. Please see Bendor and Wang (2008) for more details [23].

## Results

### Long-duration post-stimulus activity is observed in auditory cortex units during passive listening

The data analysed in this report were based on a database of 1557 single units recorded from the auditory cortex of 4 passively listening marmosets during the presentation of auditory stimuli in previous experiments. A subset of these single units exhibited poststimulus activity such as an example shown in Figure 1A. Pure tone and noise stimuli lasting 200 ms in duration were presented in order to map the rate-level function of each neuron. Pure tone stimuli were then used to find the best frequency of the neuron i.e. the tone frequency that produced the largest within-stimulus response. These pure tones were presented at 10-20 dB above threshold for neurons with monotonic rate-level functions and at preferred levels for non-monotonic neurons. This mapping approach resulted in multiple stimulus types being presented to each neuron. As quantification of post-stimulus activity was based on significant difference from baseline firing, units were required to have a spontaneous firing rate of >1 spike per second and to have been presented with 5 or more repetitions of each stimulus to be included in all further analysis. This criterion was met by 1188 units.

**Figure 1:**
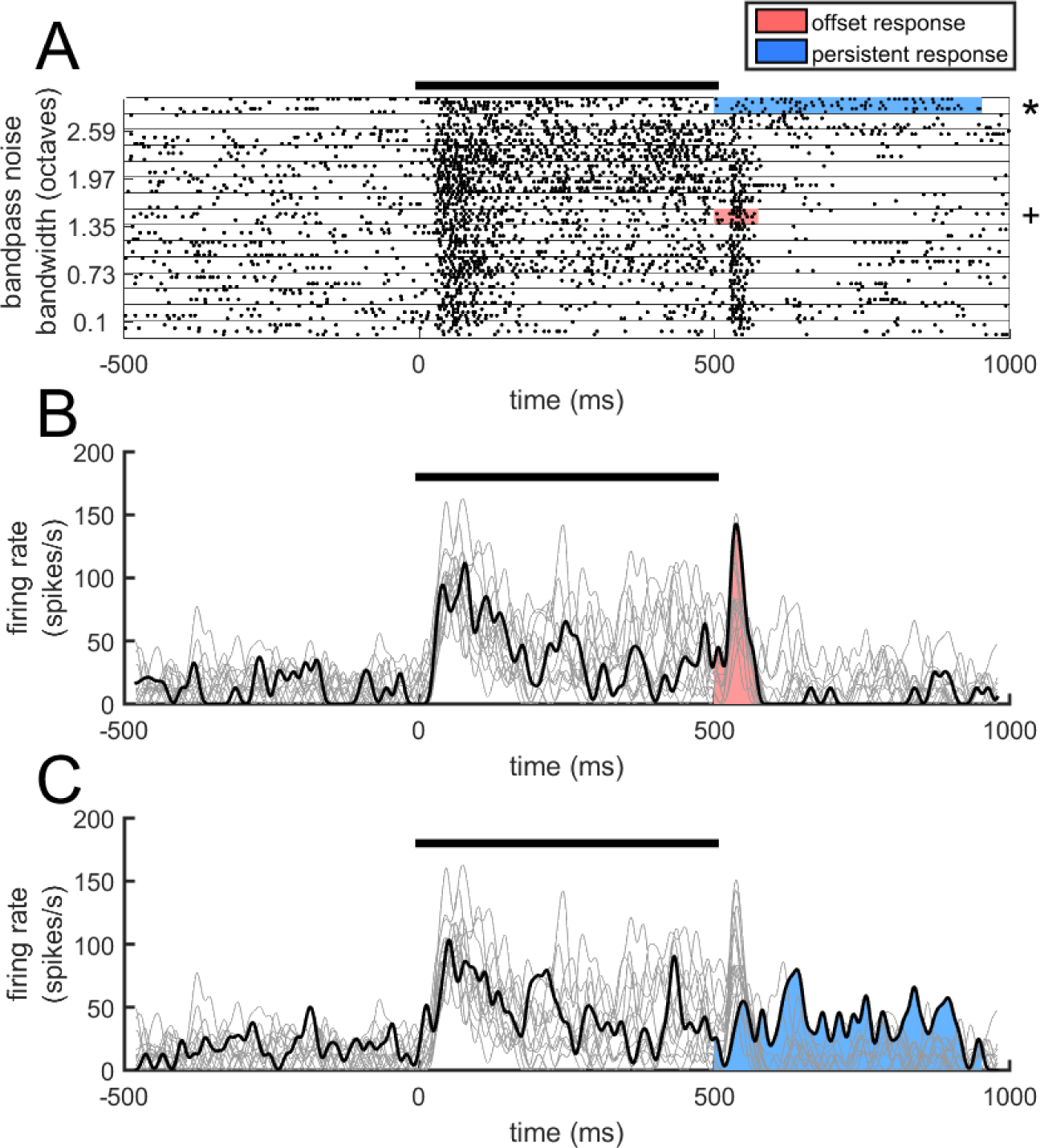
Post-stimulus activity during passive listening. A. Example unit displaying short-duration offset responses to the majority of stimuli but also long-duration post-stimulus activity in response to particular stimuli. Spike rasters showing a representative offset response are indicated by the +. Spike rasters showing a long-duration post-stimulus response are indicated by the *. The within-stimulus period is indicated by the black bar while the duration of significant post-stimulus activity on these trials in indicated by grey shading. B. The black trace corresponds to the PSTH of spiking activity indicated by the + symbol in A while the grey traces are the PSTHs associated with all other stimuli. The black bar indicates the stimulus duration. The coloured area of the PSTH indicates the duration of the post-stimulus response that was elevated above baseline. C. PSTH of spiking activity indicated by the * symbol in A, showing long-duration post-stimulus activity in the same unit.

The duration of post-stimulus activity was quantified by convolving spike trains for a given stimulus with a bi-directional Gaussian filter (σ=5ms, total bandwidth=20ms) in order to estimate the mean instantaneous firing rate before, during and after the presentation of the stimulus. Significant post-stimulus activity was defined as periods of firing that were over 2 standard deviations of baseline activity over mean baseline firing. The observed post-stimulus activity could be highly variable over time. Therefore, momentary drops below this threshold (*<*20ms) were not considered in our estimate of post-stimulus activity duration. In order to control for chance fluctuations in firing rates, the duration of elevated firing during spontaneous activity during the pre-stimulus period was also calculated. The duration of post-stimulus activity was required to be above the range of values that were observed by chance in the pre-stimulus period.

Short-duration offset responses were commonly observed, with 76.43% of units showing significant post-stimulus responses beginning within 50ms of stimulus offset and lasting *<*100ms (Figure 1B). By searching the stimulus space, it was possible to evoke post-stimulus activity meeting the above criteria in 81.4% of the units recorded (Figure 1C). Within the latter population, 61.12% of units also showed short-duration offset responses in response to at least one stimulus.

Stimulus parameters such as stimulus type and inter-stimulus interval (ISI) were varied across a subset of the neurons recorded. Subjects were presented with a variety of noise and tone stimuli with a duration of 200ms. Stimuli were separated by a 300ms inter-stimulus interval (ISI) during which post-stimulus activity duration was quantified. For the population of 560 units recorded with these stimulus parameters, the longest duration post-stimulus response observed for each unit exceeded this 300ms interval in 31.25% of units (Figure 2A). When the ISI was extended to 500ms 5.93% of the 118 units tested exceeded this 500ms interval (Figure 2B). For units presented with stimuli followed by an ISI of >500ms (N = 57), the range of post-stimulus response durations spanned from 12ms to 1681ms (Figure 2C).

**Figure 2:**
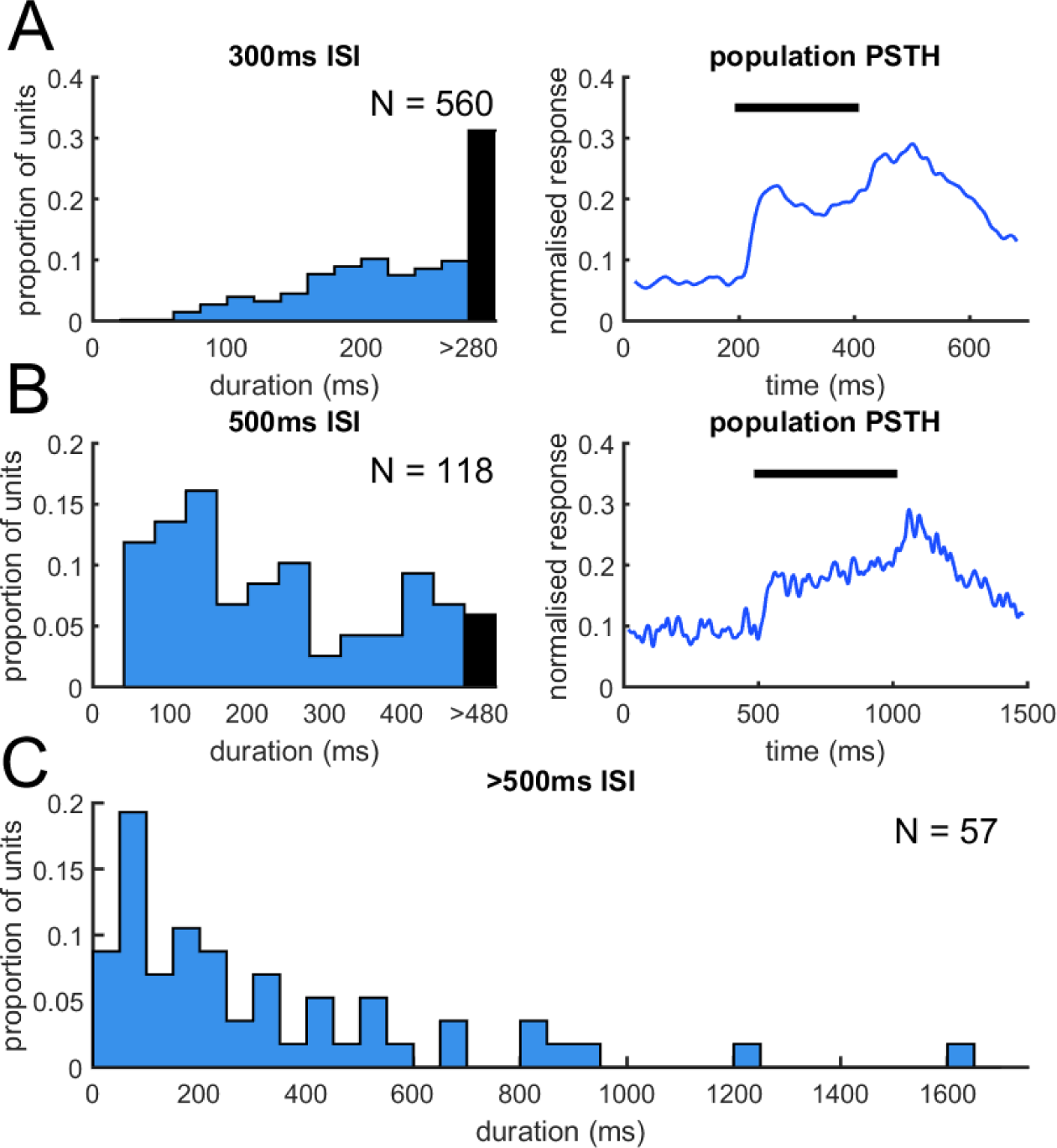
Duration of post-stimulus for different stimulation parameters. A. Left panel: Distribution of longest duration post-stimulus responses for 560 single units presented with 200ms stimuli followed by a 300ms inter-stimulus interval (ISI). Right panel: Normalised population PSTH for these responses. B. Left panel: Distribution of longest duration post-stimulus responses for 118 single units presented with 500ms stimuli followed by a 500ms ISI. Right panel: Normalised population PSTH for these responses. C. Distribution of longest duration post-stimulus responses for 57 single units presented with stimuli that were followed by an ISI >500ms.

### Post-stimulus activity is observed following adapting, sustained and suppressed responses

The response profiles observed within auditory stimulation were examined next in order to gain insight into the dynamics that might produce this post-stimulus activity. Within-stimulus evoked activity was quantified by taking the ratio of the within-stimulus firing rate and the baseline firing rate preceding sensory stimulation to produce an evoked ratio (Figure 3A). An evoked ratio >1 would indicate that the within-stimulus firing rate was increased with respect to baseline firing while values *<*1 would indicate suppression of firing in the within-stimulus period. The dynamics of responses with an evoked ratio of >1 were first analysed. For each response in this group the median within-stimulus spike time was calculated as a measure of the extent of adaptation that occurred during sensory stimulation. This median spike time was normalised by the duration of the stimulus, producing a normalised median spike time (NMST) for each unit between 1 and 0. This measure of adaptation results in fast adapting responses being associated with values near to zero and ramping responses being associated with values near one. A continuous distribution of NMSTs was observed, with the majority of units responding with sustained responses, associated with NMSTs in the middle of the range (Figure 3A). Adapting responses were defined as those with an NMST of *<*0.35 while sustained responses were defined as those with an NMST of >0.35 (Figure 3B). Long-duration post-stimulus activity was observed following both adapting and sustained responses (Figure 3C). NMSTs were also calculated for the mean response to all stimuli (unit NMST; Figure 3D) and followed a similar distribution to NMSTs calculated for responses to a single stimulus (response NMST; Figure 3B). Response NMSTs were significantly correlated with unit NMSTs (r = 0.61, p*<*0.001) indicating that within-stimulus response profiles that precede long-duration post-stimulus activity are representative of the average response dynamics of the units (Figure 3E).

**Figure 3:**
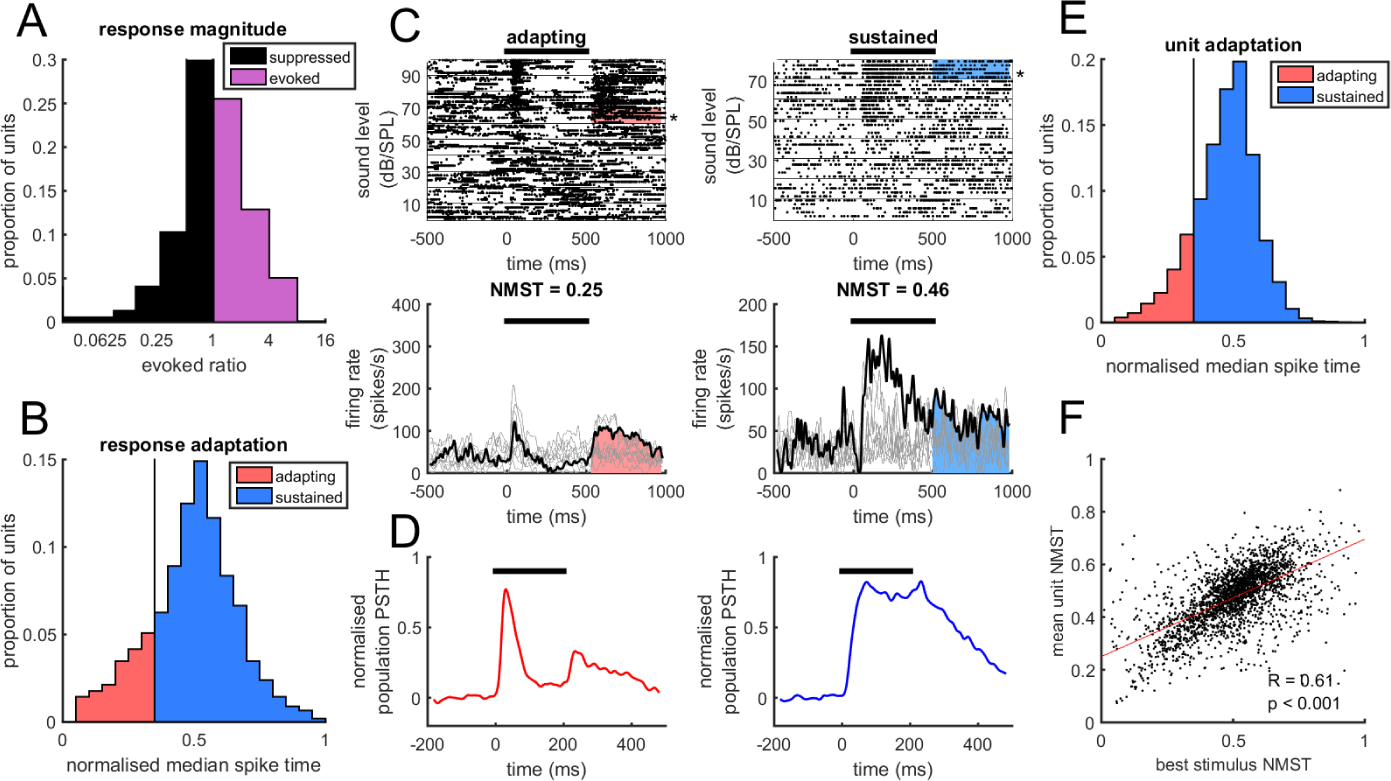
Post-stimulus activity following adapting and sustained responses. A. Distribution of evoked ratios. The vertical line at 1 indicates the threshold separating evoked from suppressed responses. B. Distribution of normalised median spike times of responses to the stimulus that evoked the longest duration post-stimulus activity. The vertical line at 0.35 indicates the threshold separating adapting from sustained responses. C. Upper panels: Raster plots and corresponding PSTHs for two units. The left panel shows an adapting response while the right panel shows a sustained response. Lower panels: Raster plots and corresponding PSTHs for the same units. PSTHs in bold indicates the responses to the stimulus that produced the longest duration post-stimulus activity for that unit. Titles above PSTHs show the normalised median spike time for this response. D. Normalised population PSTH for adapting (left panel) and sustained (right panel) responses. E. Distribution of normalised median spike times calculated from the mean responses across all stimuli for each unit. F. Scatter plot between unit and response normalised median spike times showing a positive correlation.

The relationship between within-stimulus response magnitude and the duration of post-stimulus activity was next examined. Within-stimulus tuning curves were calculated by measuring the peak firing rate during the stimulus time period in the mean PSTH for each stimulus (Figure 4A). Post-stimulus tuning curves were calculated by measuring the duration of significant post-stimulus activity produced by each stimulus. A correlation coefficient (cc) was then calculated for these tuning curve pairs for each unit by correlating the peak firing rates across stimuli during the within period with the duration of post-stimulus activity. Tuning curves showed a significant positive correlation across the population of units classified as having adapting responses to their best stimulus (median = 0.31, sign rank; p*<*0.001) (Figure 4B). This was also the case for units that showed sustained responses to their best stimulus (median = 0.34, sign rank; p*<*0.001) (Figure 4C). A significant positive correlation was observed between NMSTs and tuning curve ccs (r = 0.25, p*<*0.001) (Figure 4C), indicating that tuning curve ccs were weaker for units with lower NMSTs, i.e. adapting units, and stronger for those with higher NMSTs, i.e. sustained units. In keeping with the correlation between NMSTs and tuning curve ccs (Figure 4C), tuning curves for units that showed a sustained response showed a significantly greater correlation across the population than units that showed an adapting response to their best stimulus (Adapting median cc = 0.31, Sustained median cc = 0.34, rank-sum test; p*<*0.001).

**Figure 4:**
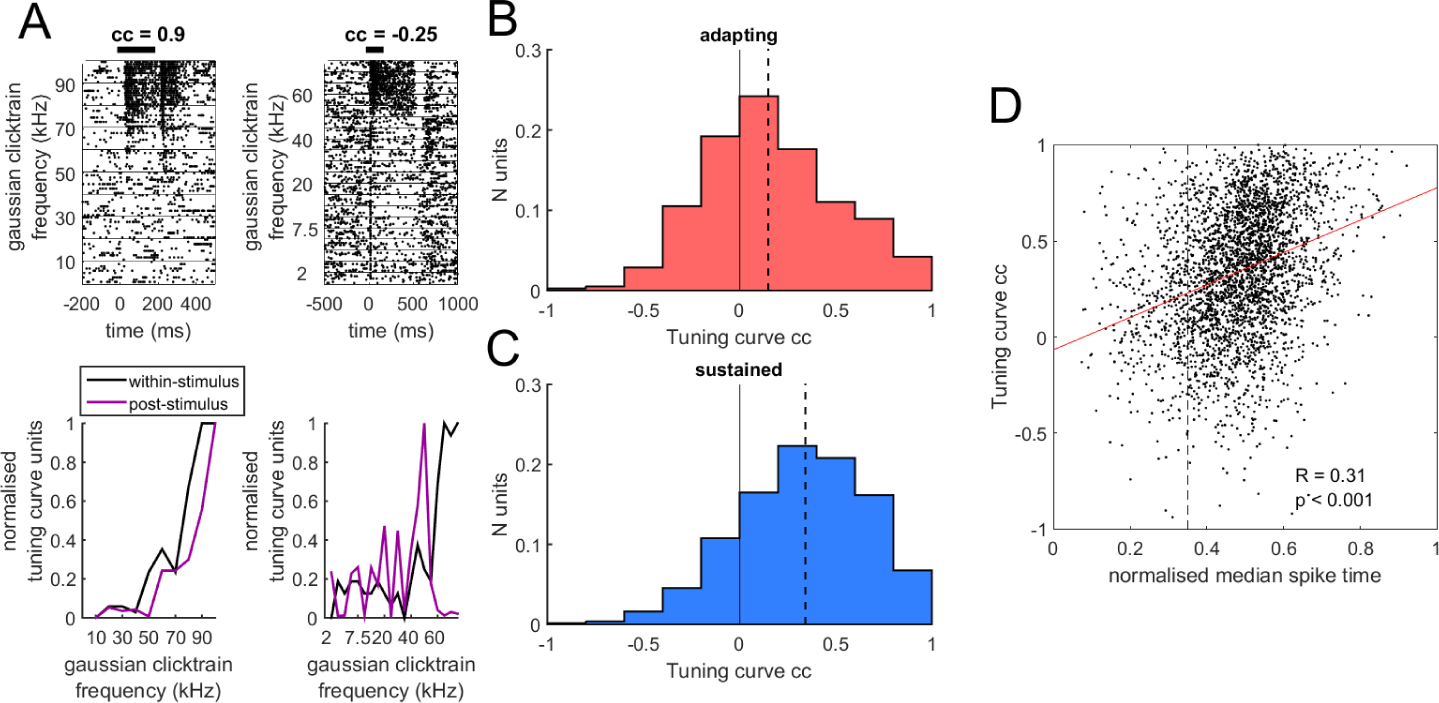
Within and post-stimulus tuning curves. A. Rasterplots of responses (upper panels) and tuning curves (lower panels) for two units. Black curves show the within-stimulus tuning curves, calculated by taking the normalised peak firing rate in response to each stimulus. The grey curves show the post-stimulus tuning curve for the unit, calculated by measuring the duration of significantly elevated firing in the inter-stimulus interval. B. The distribution of correlation coefficients (ccs) between the within and post-stimulus tuning curves for units showing adapting responses to their best stimulus. The broken line indicated the median cc for the distribution. C. Distribution of ccs between the within and post-stimulus tuning curves for units showing sustained responses to their best stimulus. D. Scatter plot of normalised median spike times (NMSTs) against tuning curve ccs, showing a positive correlation.

Not all units that showed significant post-stimulus activity showed evoked within-stimulus auditory activity in the within-stimulus interval. These units, associated with an evoked ratio of *<*1, were examined next. Units in this population typically demonstrated within-stimulus suppression (Figure 5A) and also showed long-duration post-stimulus activity (Figure 5B). For stimuli with an ISI of 300ms, significant variation in post-stimulus response duration was observed between adapting, sustained and suppressed response subtypes (Figure 5C)(ANOVA(2); p*<*0.001). Pairwise significance tests between group means (Tukey-Kramer test, corrected for multiple comparisons) indicated that sustained responses were followed by longer duration post-stimulus responses than suppressed responses (adapting vs sustained: p*<*0.001, adapting vs suppressed: p=0.77, sustained vs suppressed: p*<*0.001). The same pattern was observed for stimuli with an ISI of 500ms (ANOVA(2); p*<*0.001, adapting vs sustained: p*<*0.001, adapting vs suppressed: p=0.64, sustained vs suppressed: p*<*0.001).

**Figure 5:**
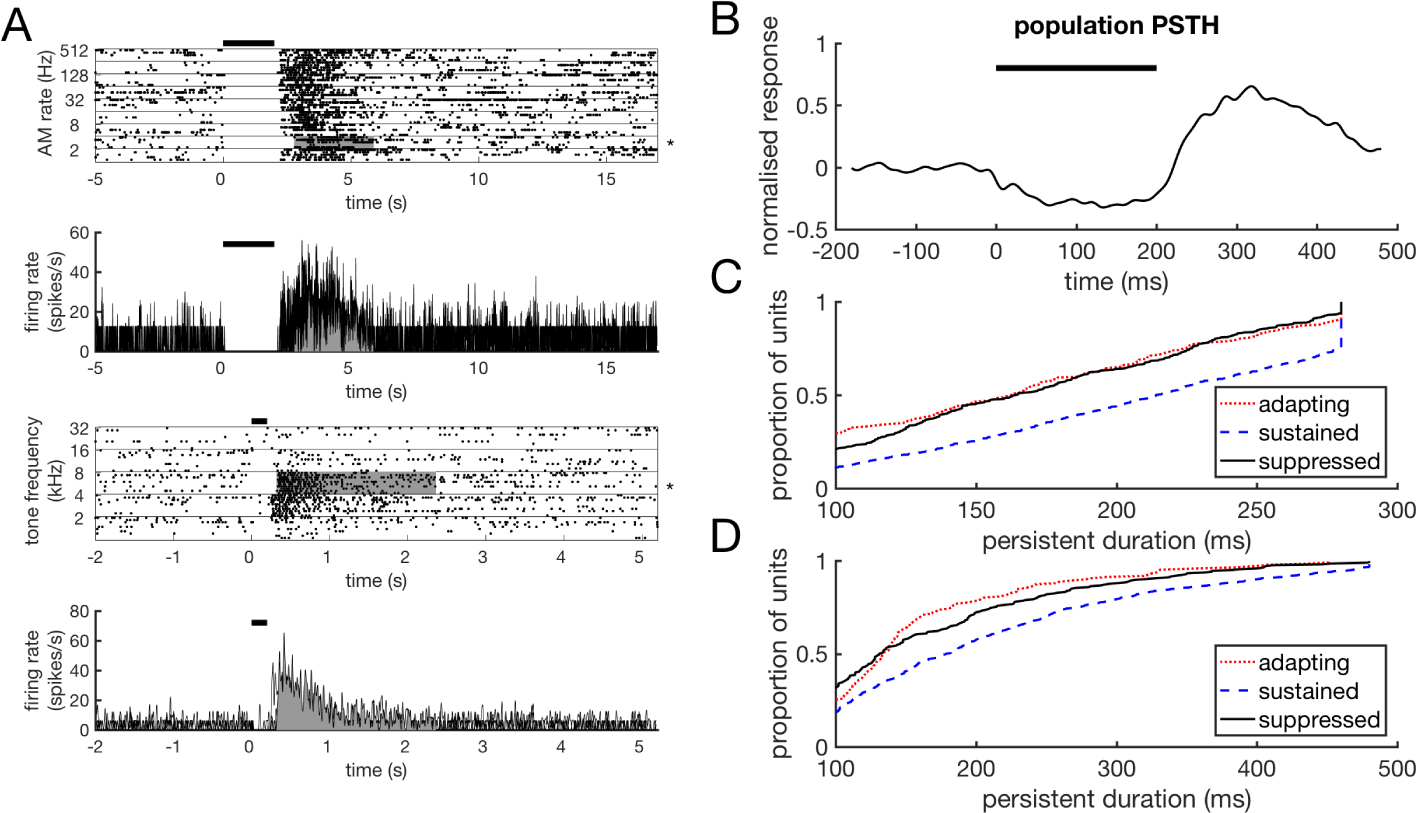
Post-stimulus activity following stimulus induced suppression. A. Raster plots and corresponding PSTHs for two suppressed units. PSTHs in bold indicate the responses to the stimulus that produced the longest duration post-stimulus activity for that unit. B. Normalised population PSTH for units showing within-stimulus period suppression. C. Cumulative distribution of longest duration post-stimulus responses for adapting, sustained and suppressed responses for stimuli with an inter-stimulus interval of 300ms. C. Cumulative distribution of post-stimulus response durations for stimuli with an inter-stimulus interval of 500ms.

A significant positive correlation was observed between baseline firing rates and the duration of post-stimulus activity across adapting (r = 0.32, p*<*0.001) (Figure 6A), sustained (r = 0.23, p*<*0.001) (Figure 6B) and suppressed units (r = 0.1, p*<*0.01) (Figure 6C). We quantified the burstiness of baseline firing using the coefficient of variation, *CV*:

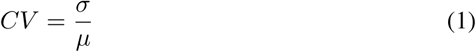

**Figure 6:**
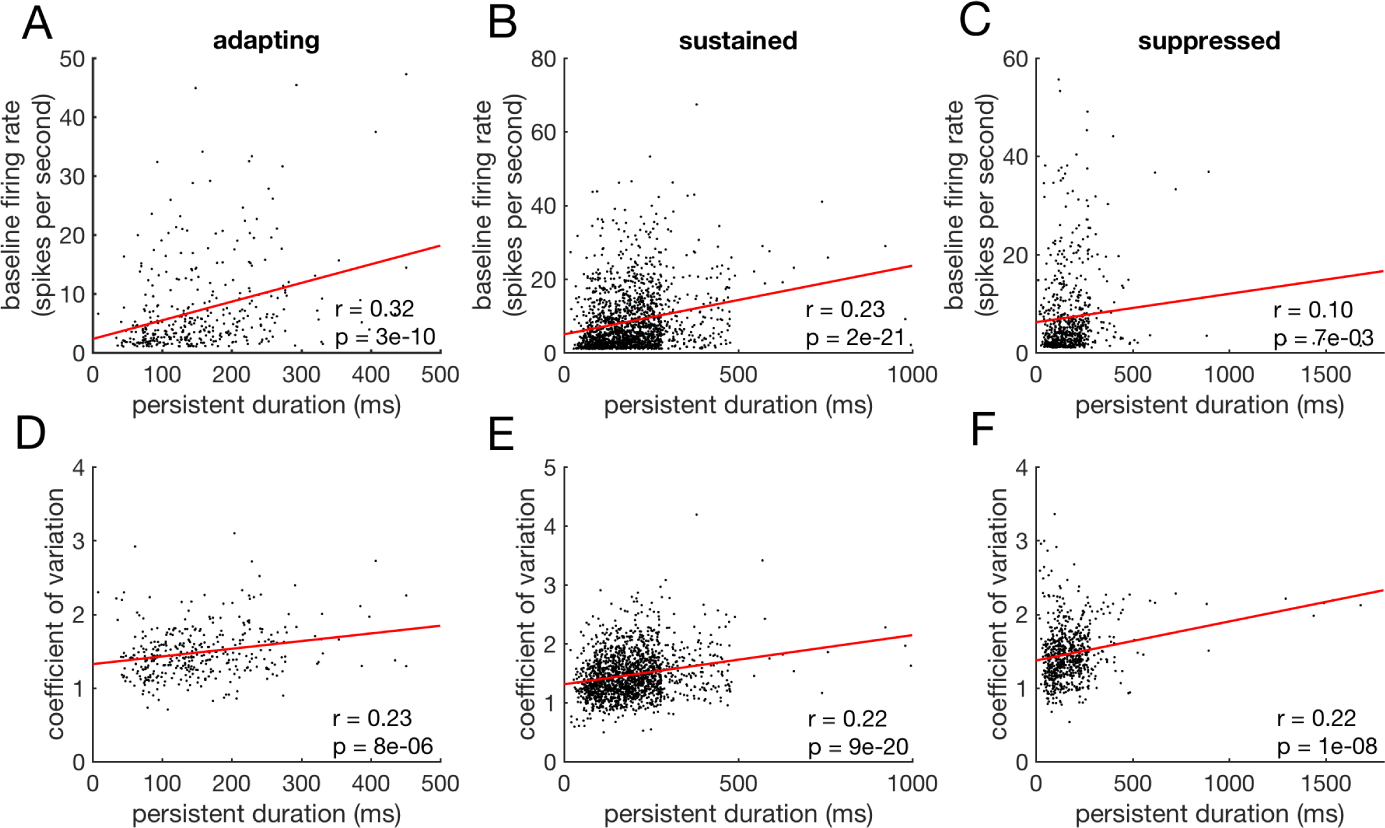
Baseline firing and burstiness is positively correlated with post-stimulus activity duration across response types. A. Correlation between baseline firing rates and the duration of post-stimulus activity for adapting units. B. Correlation between baseline firing rates and the duration of post-stimulus activity for sustained units. C. Correlation between baseline firing rates and the duration of post-stimulus activity for suppressed units. D. Correlation between the coefficient of variation and the duration of post-stimulus activity for adapting units. E. Correlation between the coefficient of variation and the duration of post-stimulus activity for sustained units. F. Correlation between the coefficient of variation and the duration of post-stimulus activity for suppressed units. Red lines indicate the regression slope for each data pair shown.

Where the standard deviation of the inter-spike-intervals is σ and the mean inter-spike-interval is *µ*. A significant positive correlation was also observed between the coefficient of variation and the duration of post-stimulus activity across adapting (r = 0.23, p*<*0.001) (Figure 6D), sustained (r = 0.22, p*<*0.001) (Figure 6E) and suppressed units (r = 0.22, p*<*0.001) (Figure 6F).

### Post-stimulus activity emerges in cortex

Single units (N = 379) were also recorded from the auditory thalamus of passively listening marmosets [26, 28] (Figure 7A). The criterion of having a baseline firing rate of >1 spikes per second was met by N = 361 units. Short-duration offset responses were also observed in the majority of these units (76.45%). It was possible to evoke post-stimulus activity exceeding the duration of spontaneous elevations in firing rate in 81.44% of thalamic units. When presented with stimuli lasting 200ms followed by a 300ms ISI (N units = 153), thalamic post-stimulus activity was significantly shorter than the post-stimulus activity reported above in cortex (rank-sum test, p*<*0.001) (Figure 7A). Thalamic post-stimulus activity was also significantly shorter than cortical activity for units presented with stimuli followed by an ISI of >300ms (N units = 102) (rank-sum test, p*<*0.001) (Figure 7B). The latency of post-stimulus activity, the time following stimulus termination before firing became significantly elevated, was significantly shorter in cortex (rank-sum test, p*<*0.001)(Figure 7C).

**Figure 7:**
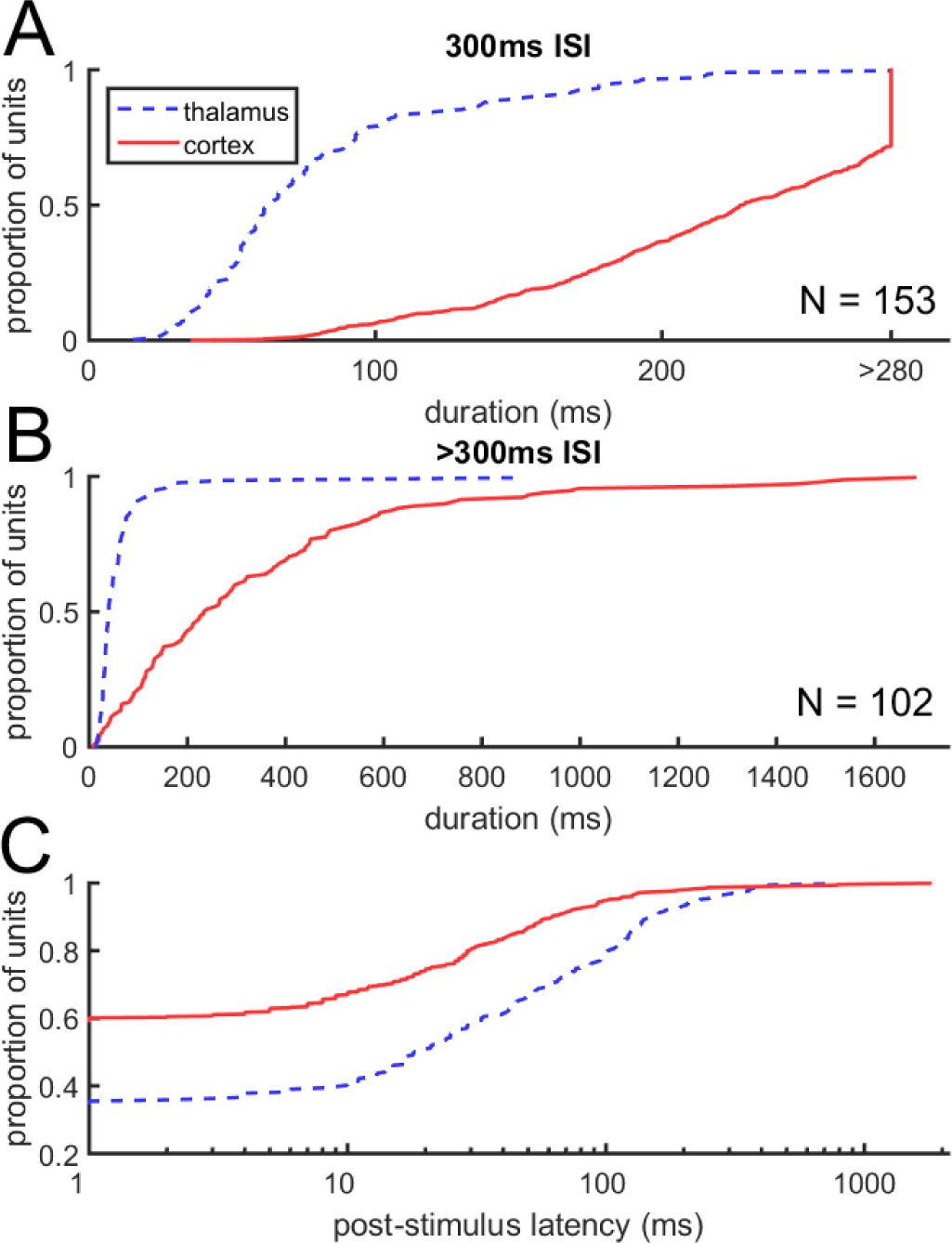
Post-stimulus activity in auditory thalamus. A. Cumulative distribution of longest duration post-stimulus responses for thalamic units (broken line) presented with 200ms stimuli followed by a 300ms inter-stimulus interval (ISI). Ns refer to thalamic units. The cumulative distribution of cortical response durations is shown for comparison (solid line). B. Cumulative distribution of longest duration post-stimulus responses for thalamic units presented with stimuli that were followed by an ISI >300ms. Again, the cumulative distribution of cortical response durations is shown for comparison (solid line). C. Cumulative distribution of post-stimulus activity latency in thalamus (broken line) and cortex (solid line).

We next investigated whether variation in the duration of post-stimulus activity exists between cortical fields. Single units were recorded in ares A1, R and RT in core auditory cortex of two subjects (Figure 8A). In order to compare durations across fields it was necessary to only include units for which the maximum duration of post-stimulus activity could be accurately estimated, this is, where the duration of activity did not exceed the inter-stimulus interval. The duration of post-stimulus activity showed significant variation across cortical fields (Kruskall-wallis test, H(2)=32.46, p*<*0.001) (Figure 8B). Post-hoc multiple comparisons of mean ranks indicated that post-stimulus activity of significantly longer duration in RT compared to A1 and R (median durations, A1=174ms, R=186.5ms, RT=214ms; A1 vs R, p=0.04; A1 vs RT, p*<*0.001; R vs RT, p=0.002). The pattern of longest duration responses being found in RT also held across individual subjects (Subject 1: Kruskall-wallis test, H(2) = 10.86, p=0.004; median durations, A1=150ms, R=167ms, RT=187ms; A1 vs R, p=0.065; A1 vs RT, p=0.008; R vs RT, p=0.536; Subject 2: Kruskall-wallis test, H(2)=23.74, p*<*0.001; median durations, A1=169ms, R=162ms, RT=219ms; A1 vs R, p = 0.531; A1 vs RT, p*<*0.001; R vs RT, p*<*0.001). Finally, we investigated whether the proportion of units showing each of the three within-stimulus response profiles (adapting, sustained, suppressed) varied across cortical fields. In A1 the majority of units that showed post-stimulus activity showed sustained responses in the within-stimulus period (57.3%; Figure 8C). The proportion of units showing sustained responses was even greater in R (70.6%) and greater still in RT (75.1%). This indicates that response profiles preceding post-stimulus activity become more sustained in the more anterior auditory cortical fields.

**Figure 8:**
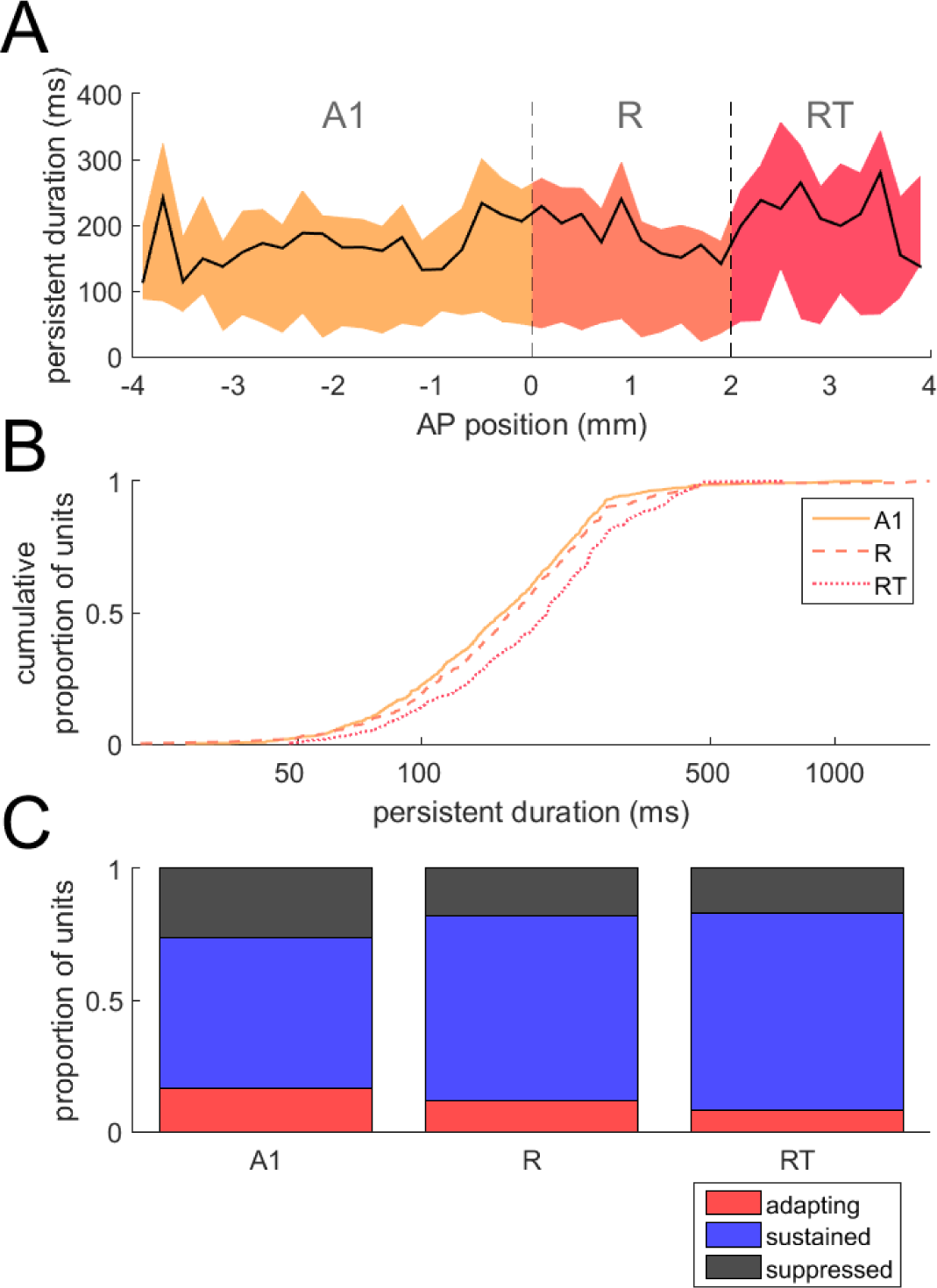
Variation in post-stimulus activity duration across auditory cortical fields. A. Median (solid line) and inter-quartile ranges (coloured areas) of post-stimulus activity across the anterior-posterior extent of auditory cortex. Boundaries of cortical fields A1, R and RT are indicated with dashed lines. Data shown is combined from two subjects where recordings were made across the entire anterior-posterior extent of auditory cortex. B. Cumulative distributions of post-stimulus activity across cortical fields showing a significant increase in duration in more anterior fields, R and RT. C. Proportions of response types observed in different cortical fields, showing a trend towards a greater proportion of sustained responses in more anterior fields.

## Discussion

Persistent responses in auditory cortex following the termination of an auditory stimulus have only previously been reported during performance of tasks requiring working memory [17]. Here we report the presence of long-duration post-stimulus responses in a population of auditory cortical neurons in the absence of a task. Single units were recorded in the auditory cortex (A1, R and RT) and thalamus of awake, passively-listening marmosets. Initially, we were only able to measure post-stimulus responses up to 300ms given our stimulus parameters. We found a significant proportion of units (31.25%) showed significantly elevated firing for this entire duration in response to tone and noise stimuli. When this interval was extended to 500ms it was possible to accurately measure the duration of this activity in of the majority of units, although in 5.93% of units post-stimulus responses exceeded the full length of this interval. When the interval was extended beyond this we found a minority of units that not only exceeded 500ms but showed persistent activity exceeding one second. The number of units that continue to be persistently active across a post-stimulus period therefore appears to decrease over time, in keeping with the reports of population responses in auditory cortex becoming increasingly sparse during sensory stimulation [29].

Post-stimulus activity was observed following several different response profiles during sensory stimulation. As previously reported, units in auditory cortex are capable of showing adapting and sustained responses [30]. The evoked responses of units fell along a continuum from adapting, through sustained to ramping. Long-duration post-stimulus activity was observed following both adapting and sustained response profiles. We also found that the extent of adaptation on trials that produced post-stimulus responses was representative of the mean response profile of the units to a variety of stimuli. This allowed us to examine the relationship between within-stimulus and post-stimulus tuning for adapting and sustained response types. For sustained responses, the peak firing rate in the within stimulus period predicted the duration of post-stimulus activity; however, this effect was weaker for adapting units. This indicates that these response types might arise from different mechanisms.

One likely candidate mechanism for post-stimulus activity following adapting responses is a difference in the balance of excitation and inhibition. Inhibition is known to play a role in the generation of adapting onset responses [31] and thus may be responsible for altering the tuning of the post-stimulus responses. For post-stimulus activity following sustained responses, however, there is little evidence for an inhibitory mechanism. Cortical networks have a highly recurrent architecture [32] which may play a role in the generation and maintenance of these responses through an entirely excitatory mechanism. A recurrent mechanism of this kind would account for the temporal fluctuations in post-stimulus activity that were widely observed in these units. The longer duration of post-stimulus activity following sustained responses is also in keeping with a mechanism in which excitation dominates over inhibition during the post-stimulus period. We also observed units that showed dramatic suppression in response to auditory stimuli, followed by the longest duration post-stimulus activity that we observed. This dramatic suppression is in keeping with a role for inhibition in generating this post-stimulus activity. Adaptation of inhibitory signals during sensory stimulation may result in a period of post-stimulus dominance for excitation in these units as well as in the adapting units. A post-stimulus imbalance in excitatory and inhibitory balance may therefore provide a common mechanism for post-stimulus activity during passive listening. Causal manipulations of inhibitory tone and recurrent activity will be required in order to parse the contributions of these different mechanisms to the generation of different forms of post-stimulus activity.

Finally we examined whether any variation in the duration of these responses exists throughout the thalamocortical system. We found that post-stimulus responses were of shorter duration in auditory thalamus compared to cortex. Post-stimulus activity was also found to have a longer latency following stimulus offset in thalamus compared to cortex. This indicates that units displaying such activity in thalamus may simply inherit this activity via cortical feedback projections. Taken together, these findings indicate that post-stimulus activity is generated in auditory cortical networks. Rebound calcium burst activity lasting hundred of milliseconds has been observed in the auditory thalamus [33] and may contribute to the emergence of the post-stimulus activity observed in this area. This mechanism may also contribute to the generation of post-stimulus activity in adapting and suppressed cortical units as rebound activity is produced following inhibitory activity [34].

Long-duration post-stimulus activity was observed in all cortical fields tested. This finding is in keeping with a model in which the circuit architecture required to generate these responses is intrinsic to cortex, resulting in the presence of these responses across cortical fields. It is not known whether such a mechanism would generalise to the cortices of other species, which show some differences in architecture [35, 36]. Differences in the organisation of inhibitory circuitry may be one potential source in interspecies variation in post-stimulus activity [37]. Persistent activity has also been observed in rodents however [38], indicating that rodent cortex is capable of generating this form of activity in the absence of a stimulus. By systematically searching stimulus space it may therefore be possible to evoke such activity during passive listening in rodent cortex.

The function of post-stimulus activity has been studied in a variety of behavioural contexts but the data presented here indicate that this activity may also play a role during passive listening. Echoic memory is a form of short-term sensory memory that differs from working memory in that it is posited to be active under all behavioral conditions, including passive listening [39]. The content of echoic memory is generally stored on the order of seconds [40, 41, 42], a timescale that fits with the duration of post-stimulus activity observed here. An alternate possibility is that the activity reported here reflects the recruitment of circuits that exist in order to retain information in working memory and provide no function during passive listening.

A functional role for this activity appears plausible, however, given the requirement of temporal integration in auditory perception in all species, from insects [43] to humans [44]. Investigating the function of activity observed in the absence of a behavioural task is particularly challenging however. Experiments addressing the function of this activity could exploit behavioural measures that provide an implicit measure of the processing of sensory information in the absence of a task, such as fear-conditioned freezing. Combining this approach with temporally precise suppression of this activity through optogenetic methods, it may be possible to investigate the potential role of this activity in echoic memory and temporal integration.

The presence of sustained auditory cortical activity long after the termination of a stimulus is potentially problematic for models of perception in which there is a necessary correspondence between activity in cortex and the occurrence of a percept [45]. It appears possible that the sparse nature of this signal in the cortical population may provide a distinct signature that can allow it be be differentiated from perceptual signals by downstream brain areas. Taken in conjunction with the presence of spontaneous upstates in sensory cortex [46], however, the presence of this activity provides evidence that the subjective percept of a sensory stimulus is likely not simply generated as a necessary result of the occurrence of cortical activity.

